# Light quantity impacts early response to cold and cold acclimation in young leaves of Arabidopsis

**DOI:** 10.1101/2024.11.27.625718

**Authors:** Markéta Luklová, Marieke Dubois, Michaela Kameniarová, Klára Plačková, Jan Novák, Romana Kopecká, Michal Karady, Jaroslav Pavlů, Jan Skalák, Sunita Jindal, Ljiljana Tubić, Zainab Quddos, Ondřej Novák, Dirk Inzé, Martin Černý

**Affiliations:** Department of Molecular Biology and Radiobiology, Faculty of AgriSciences, Mendel University in Brno, 61300 Brno, Czech Republic; Department of Plant Biotechnology and Bioinformatics, Ghent University, 9000 Ghent, Belgium; VIB Center for Plant Systems Biology, 9000 Ghent, Belgium; Laboratory of Growth Regulators, Institute of Experimental Botany, The Czech Academy of Sciences & Palacký University, Šlechtitelů 27, CZ-779 00 Olomouc, Czech Republic; Present address: Department of Plant and Crops, Faculty of Bioscience Engineering, Ghent University, Ghent, Belgium; Department of Botany and Zoology, Faculty of Science, Masaryk University, Kotlarska 2, 611 37 Brno, Czech Republic; Present address: CEITEC – Central European Institute of Technology, Masaryk University, CZ-62500, Brno, Czech Republic; Present address: Biology Centre CAS, Branišovská 1160/31 370 05 České Budějovice, Czech Republic; Institute for Biological Research “Siniša Stanković”, National Institute of the Republic of Serbia, University of Belgrade, Bulevar despota Stefana 142, 111 08 Belgrade, Republic of Serbia

**Keywords:** acclimation, freezing tolerance, transcriptome, proteome, lipidome, jasmonic acid, leaf development

## Abstract

Plant reactions to stress vary with development stage and fitness. This study assessed the relationship between light and chilling stress in Arabidopsis acclimation. By analyzing the transcriptome and proteome responses of expanding leaves subjected to varying light intensity and cold, 2251 and 2064 early response genes and proteins were identified, respectively. Many of these represent as yet unknown part of early response to cold, illustrating development-dependent response to stress and a duality in plant adaptations. While standard light promoted photosynthetic upregulation, plastid maintenance, and increased resilience, low light triggered a unique metabolic shift, prioritizing ribosome biogenesis and lipid metabolism and attenuating expression of genes associated with plant immunity. The comparison of early response in young leaves with that in expanded ones showed striking differences, suggesting a sacrifice of expanded leaves to support young ones. Validations of selected DEGs in mutant background confirmed a role of HSP90-1, transcription factor FLZ13, and Phospholipase A1 (PLIP) in response to cold, and the PLIP family emerged as crucial in promoting acclimation and freezing stress tolerance. The findings highlight the dynamic mechanisms that enable plants to adapt to challenging environments and pave the way for the development of genetically modified crops with enhanced freezing tolerance.

## 1 INTRODUCTION

Given their sessile nature, plants are consistently exposed to a myriad of environmental stressors, seldom encountering singular abiotic factors (Kopecká et al., 2023). Recent studies have highlighted that simultaneous presence of environmental stresses triggers distinct molecular responses, transcending the mere aggregation of individual stress reactions (Zandalinas *et al*. 2020). The interplay between different signaling pathways can lead to a phenomenon known as acclimation. During evolution, plants developed this mechanism to increase their tolerance to abiotic stresses and to withstand the harsh conditions of the environment. The transcriptomic-based induction of the acclimation can be delineated into several distinct stages. The initial stage, occurring within seconds, involves the rapid expression of genes to prevent irreversible damage. That is followed by the activation of early response genes, which lay the foundation for long-term protection. Next, late response genes are triggered, initiating systemic acclimation mechanisms. Finally, the de-acclimation process prepares the plant for stress resolution. A disruption in any of the acclimation steps may compromise the plant’s ability to adapt and significantly impact its resilience (Zandalinas, Sengupta, Burks, Azad & Mittler 2019).

The plant’s reaction varies based on its stage of development and overall fitness (Peck and Mittler, 2020). For instance, when exposed to high-light stress, young leaves activate protective mechanisms like non-photochemical quenching, which might not be present or as effective in mature leaves (Rankenberg et al., 2021). Moreover, when subjected to stress, younger leaves tend to show increased levels of anthocyanin and possess a higher ability to remove reactive oxygen species (Zhu *et al*. 2018). Mature leaves seem to undergo a reduction in their ability to start the detoxification process, resulting in decreased tolerance to photoinhibition. The phenomenon of young leaves being prioritized for recovery has been found across various abiotic stresses. For instance, during drought stress, the accumulation of abscisic acid (ABA) initiates a process leading to the transport of carbohydrates and early senescence in mature leaves. Simultaneously, this stimulus suppresses growth and promotes the absorption of nutrients by young leaves (Bielczynski, Łaçki, Hoefnagels, Gambin & Croce 2017).

In temperate regions, sudden temperature drops and freezing stress pose significant threats to plant survival and agricultural production (Kopecká, Kameniarová, Černý, Brzobohatý & Novák 2023). Cold stress disrupts various physiological processes essential for plant growth and survival, including membrane fluidity, nutrient uptake, and energy production. Despite our knowledge of integral components of cold perception pathway, the exact mechanism has not been identified (Kerbler and Wigge, 2023). It is believed that plasma membrane is the primary source of signaling and several membrane-associated proteins have been proposed as candidate thermosensors, including ANNEXIN1 that mediates cold-triggered Ca^2+^ influx and freezing tolerance in *Arabidopsis thaliana* (Wei et al., 2021), protein COLD1 that mediates chilling tolerance in *Oryza sativa* through G-protein signaling (Ma et al., 2015), and calcium/calmodulin-regulated receptor-like kinase that modulates cold acclimation through MAP kinase cascade (Yang et al., 2010). Substantial evidence also implicates a role of the circadian clock components ELF3, LHY, and PPR7 (Jung et al. 2020; Wu et al. 2024; Kim, Kim & Somers 2024) and light perception pathway, including Phytochome B, Phytochrome-interacting factors, and interactions with chromatin (reviewed in Kerbler and Wigge, 2023).

Following the first exposure to cold shock, rapid alterations in the plasma membrane, are initiated, when diacylglycerol kinase (DGK) undergoes activation after exposure to cold temperatures, leading to the conversion of diacylglycerol (DAG) into phosphatidic acid (PA) (Arisz *et al*. 2013). This enzymatic reaction is followed by alterations in the membrane fluidity, leading to the activation of many second messengers, including ROS, inositol phosphates, and calcium ions, that are recognized by specific protein sensors (Wei *et al*. 2021). The MAP kinase cascade facilitates the regulatory phosphorylation of downstream signaling components in response to cold stress. In *Arabidopsis*, the activation of the cascade is initiated by mitogen-activated protein kinase kinase kinase (ANP1) and leads to positive regulation of freezing tolerance (Zhang and Sonnewald, 2017). The MAP kinase cascade is also responsible for regulating the activity of ROS-scavenging enzymes in order to maintain redox equilibrium during periods of cold stress. All these mechanisms lead to the activation of the main established cascade, ICE-CBF (Inducer of CBF expression, C-repeat/dehydration-responsive element-binding factor) (Chinnusamy, Zhu and Zhu, 2007; Wang et al., 2017; Hwarari et al., 2022). Multiple positive and negative regulators have been found for various transcription factors involved in cold response. These regulators include calmodulin-responsive transcriptional 3 (CAMTA3, At2g22300; Doherty et al., 2009), open stomata 1 (OST1, AT4G33950; Ding et al., 2015)), E3 ubiquitin-protein ligase HOS1 (HOS1, At2g39810; (Ishitani, Xiong, Lee, Stevenson & Zhu 1998)), E3 SUMO-protein ligase SIZ1 and 2 (SIZ1; Miura and Ohta, 2010)), inducer of CBF expression 1 (ICE1, At3g26744; Chinnusamy et al., 2003)), Transcription factor MYB15 (MYB15, At3g23250; Agarwal et al., 2006) and STRUBBELIG-receptor family 6 (SRF6, At1g53730; Knight and Knight, 2000).

The whole mechanism is quite complex, with more than 3000 identified cold-responsive genes and overlaying regulator circuits that integrate inputs from other signaling pathways, including that of phytohormones and light. For instance, positive regulator of ethylene signaling EIN3 exerts a negative regulatory effect on the expression of CBFs (Shi *et al*. 2012), and inhibitors of the jasmonic acid signaling (JAZ1 and JAZ4) interact with ICE1 and ICE2 transcription factors inhibiting their activity (Yang et al., 2019). An additional part of the regulatory circuit is thioredoxinTRX-H2, that reduces CBFs in the nucleus, ultimately activating COR genes (Lee *et al*. 2021). Light quality and quantity is critical for cold acclimation, as evidenced in recent publications (Prerostova *et al*. 2021; Novák *et al*. 2021; Kameniarová *et al*. 2022; Sugita, Takahashi, Uemura & Kawamura 2024). Mutants in *PHYB* showed upregulated CBF expression (Jiang et al., 2020) and modulated circadian clock and ROS metabolism (Luklová *et al*. 2022). In our previous work, we established a model experiment that allowed us to follow the impact of light intensity on freezing resilience (Novák *et al*. 2021) We showed that the acclimation under lower photosynthetic photon flux density (PPFD) has a contrasting mechanism to that found under standard PPFD. In our previous research, analyzing the entire plant led to a notable bias. The interpretation was predominantly based on molecular profile of fully expanded leaves, which constitute the bulk of the plant’s biomass. Here, we present a more detailed analysis focused on young leaves, the tissue known to be actively protected by the plant. Consequently, this tissue is expected to employ distinct mechanisms compared to those found in older leaves, and provide more insight into the mechanisms behind cold resilience.

## 2 RESULTS

### 2.1 Light intensity impacts cold acclimation and freezing stress resilience

Our previous work showed that both quality and quantity of light impact response to cold stress (Novák et al., 2021; Kameniarová et al., 2022). Here, to validate the previous observations made under artificial conditions of hydroponic culture, Arabidopsis plants (Col-0) were grown as described in materials and methods, and the response to freezing was monitored by calculating LT_50_ using a survival assay. The results confirmed the previous observations that low PPFD negatively impacts response to cold and validated that the results of hydroponically grown Arabidopsis are transferrable to standard cultivation. Our previous analyses targeted conditions that resulted in approximately 50 % mortality rates of control plants. Here, a more in-depth temperature profile was analyzed, and freezing similar to natural conditions was applied **(Figure 1a)**. The comparison of survival rates between standard PPFD (C) and low PPFD (CLL) at 4 °C showed statistically significant differences (p<0.001) with corresponding LT_50_ of -11.3 °C and –7.8 °C for C and CLL, respectively (**Figure 1b**). Besides the differences in the LT_50_ values, our observations indicated a development-dependent response to cold that seemed to promote the survival of younger leaves (**Figure 1c**).

**Figure 1.**
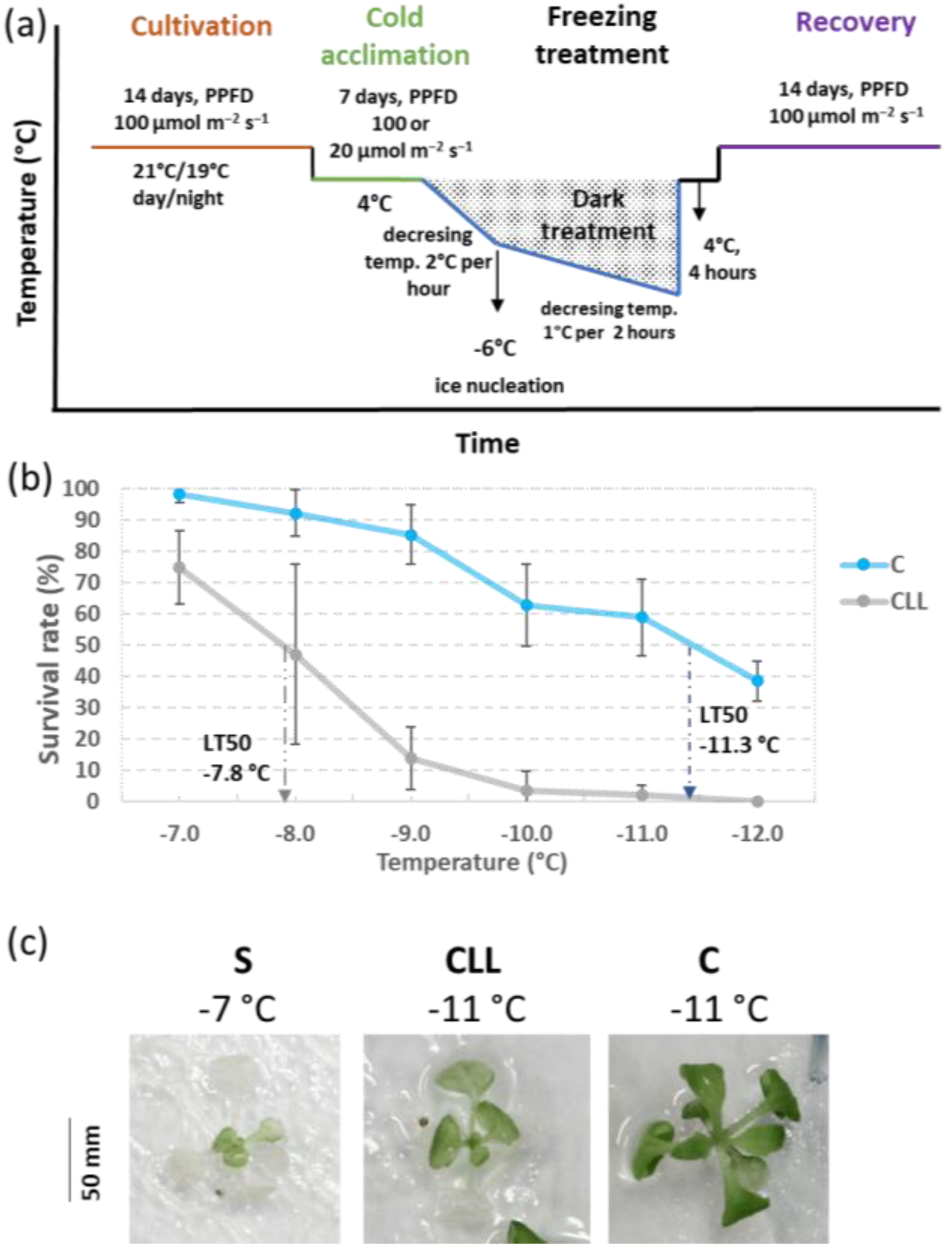
Freezing resistance in Arabidopsis plants. (a) Experimental design; (b) Freezing survival monitoring after 2 weeks of recovery period. The plot represents the means and standard deviation (three biological replicates, n>25); arrows mark calculated LT_50;_ (c) The impact of freezing stress on leaves. Representative images demonstrating the promoted survival of younger leaves documented in the survival assay experiments. *Arabidopsis thaliana* Col-0 plants were acclimated under indicated conditions (S - 100 µmol.m^-2^.s^-1^, 21°C; C - 100 µmol.m^-2^.s^-1^ 4°C; CLL - 20 µmol.m^-2^.s^-1^, 4°C) and exposed to freezing stress for two hours. The images were taken after two weeks of recovery.

### 2.2 Young leaf transcriptome in early response to cold showed significant impact of light intensity

In order to find molecular evidence for this hypothesis, the early transcriptional response to cold was analyzed in young leaves, specifically leaf n.6, which has not finished the proliferation stage. Plantlets were exposed for 3 hours to four contrasting conditions **(Figure 2)**, including (i) standard PPFD at 21°C, 100 µmol.m^-2^.s^-1^ (control; S); (ii) low-PPFD at 21°C, 20 µmol.m^-2^.s^-1^ (LL); (iii) standard PPFD at 4°C, 100 µmol.m^-2^.s^-1^ (C); (iv) low-PPFD at 4°C, 20 µmol.m^-2^.s^-1^ (CLL). A young leaves L(1.06) were collected (n=10, three fully independent biological replicates), and the transcriptome was analyzed using RNA-seq (**Figure 3a-f**, **Supplementary Tables S1**) as described in materials and methods. In parallel, the uniformity of the targeted leaves was confirmed by the cell analyses, indicating that all collected leaves were in a similar developmental stage.

**Figure 2:**
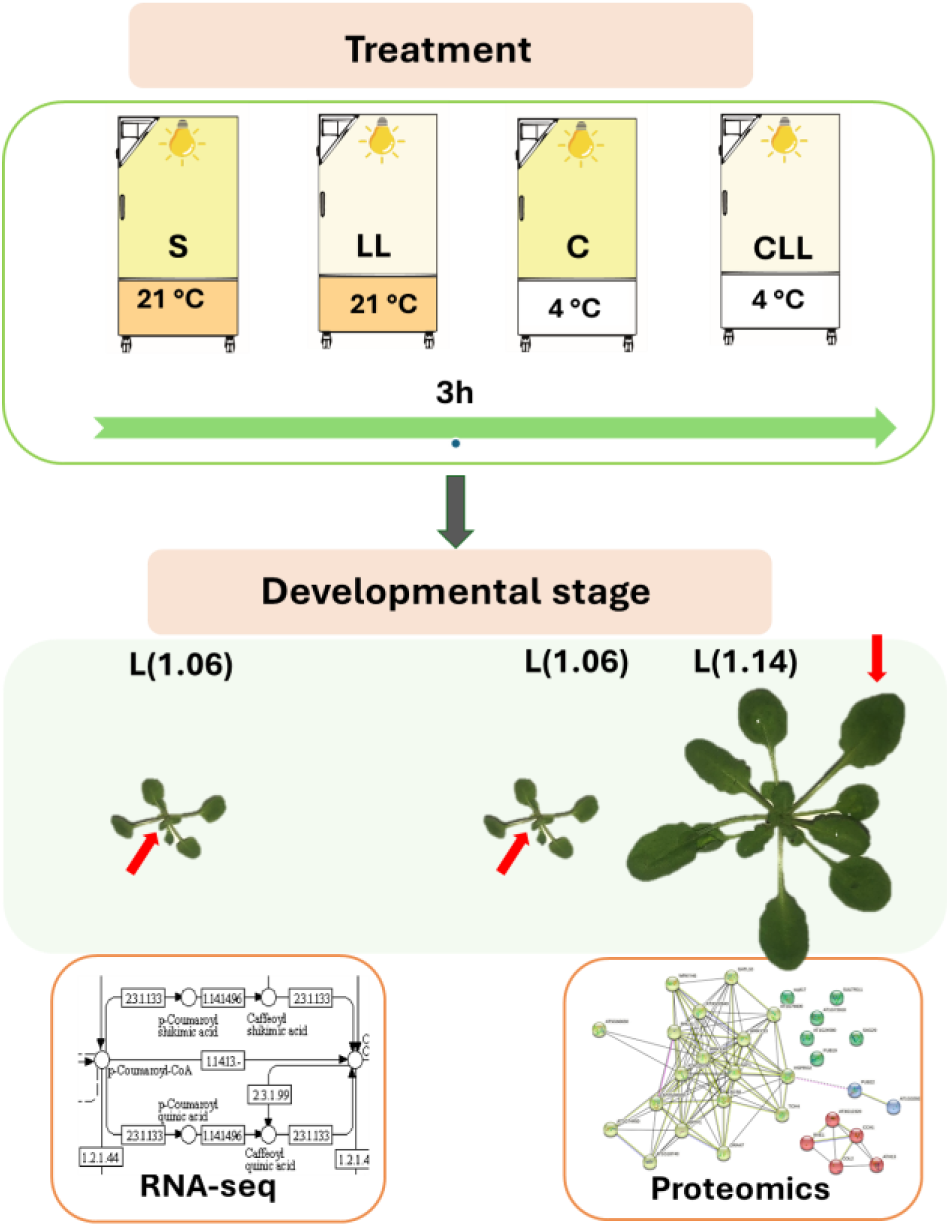
Schematic representation of experimental set-up. *Arabidopsis thaliana* plants were cultivated at standard PPFD until they reached developmental stage L1.06 or L1.14. Next, the plantlets were exposed for three hours to the following four conditions: [S] 100 µmol.m^-2^.s^-1^, 21°C; [LL] 20 µmol.m^-2^.s^-1^, 21°C; [C] 100 µmol.m^-2^.s^-1^, 4°C; [CLL] 20 µmol.m^-2^.s^-1^, 4°C. Leaves n.6 (red arrows) were collected for transcriptomics and proteomic analyses.

**Figure 3:**
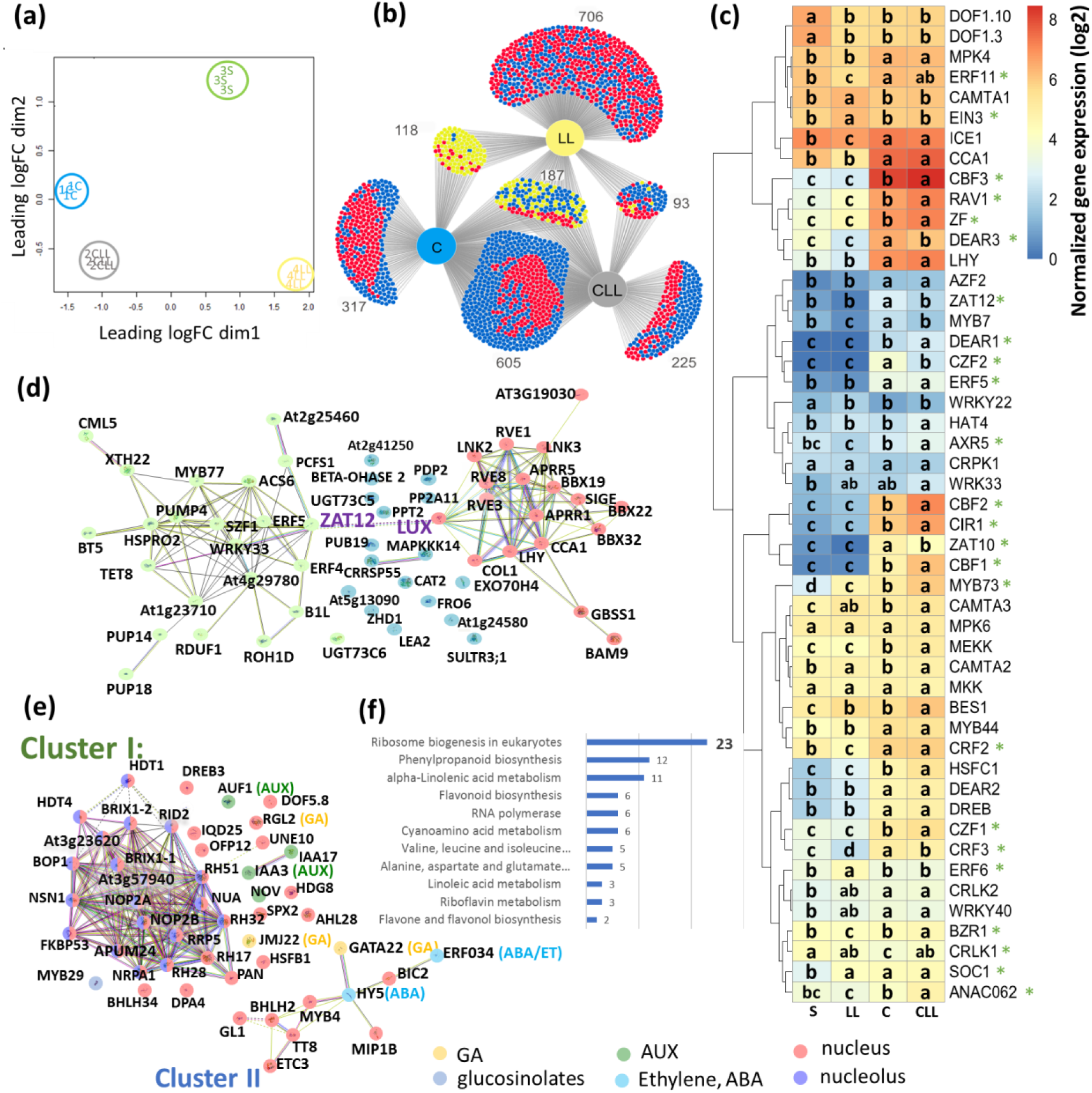
Planťs early response to cold and light treatment changes the transcriptome of the young leaf. **(a)** Multidimensional scaling of identified genes in RNA-seq analysis of young leaf L1.06; **(b)** Comparison of identified DEGs found in at least one treatment (FDR ≤ 0.05, absolute fold change >2) visualized by DiVenn (Sun *et al*. 2019). Blue and red nodes denote downregulated and upregulated genes between different treatments, respectively. Yellow nodes denote upregulation in one sample but downregulation in another; **(c)** Selected COR genes reportedly involved in early response to cold (Park *et al*. 2015; Liu, Dang, Liu, reports & 2019 2019). Significant interactions between light and cold are highlighted with an asterisk (two-way ANOVA and Tukey post hoc test, p-value ≤ 0.05); **(d)** Identified DEGs that are components of circadian clock (red dots) and circadian responsive DEGs with predicted interactions (yellow dots), and missing interacting partners (blue dots); **(e, f)** Low PPFD-specific DEGs enriched in ribosome biosynthesis, signaling, and glucosinolate metabolism and significant enrichment in metabolic pathways (p<0.05). The interactions and visualization of functional clusters were determined by STRING(Szklarczyk *et al*. 2019), minimum required interaction score = 0.4; S - 100 µmol.m^-2^.s^-1^, 21 °C; LL - 20 µmol.m^-2^.s^-1^, 21 °C; C – 100 µmol.m^-2^.s^-1^, 4 °C; CLL - 20 µmol.m^-2^.s^-1^, 4 °C. For details, see **Supplementary Table S1**.

Altogether, 28 757 genes were identified and 16 549 of these passed the set criteria (present in all replicates of at least one treatment, **Supplementary Table S1**). Multidimensional scaling of identified genes showed a clear separation of all treatments (**Figure 3a**), with C and CLL treatments being separated from LL and S in the first dimension. Next, the impact of limited light (LL), cold (C), and the combination of both factors (CLL) were analyzed in detail. The comparison of each of the three treatments with S revealed 2251 DEGs in total (relative FC > 2, FDR ≤ 0.05) and 1545 of these were cold-responsive (in C and CLL) (**Figure 3b**). The comparison of CLL and C subsets showed a significant overlap, representing 65 and 71% of identified DEGs in C and CLL, respectively. Most of these DEGs had a similar response to cold, indicating their putative role in cold response, but the degree of their respective responses differed. Next, the dataset was searched for known early cold response (COR) genes (3414 genes, Shi et al., 2017). Of these, 709 were regulated in young leaves (**Supplementary Table S1**), and many displayed a significant interaction of light and cold, as illustrated with the set of 49 early cold response genes (2-way ANOVA, p < 0.05; **Figure 3c**, **Supplementary Table S1**), including dehydration-responsive element-binding proteins CBFs (CBF1, CBF2, CBF3; **Figure 3c**).

### 2.3 Distinct C and CLL pathways: Impact on photosynthesis and phytohormone signaling and metabolism

As demonstrated previously (Novák *et al*. 2021), the CLL treatment, a combination of low-light intensity and cold induced cold acclimation process through different mechanisms to that of C. Here, 317 DEGs (**Figure 3b)** were found only in C plants. GO analysis highlighted impact on photosynthesis (FDR = 9.9e-05), photosynthesis-light reaction (FDR = 0.0041), protein-chromophore linkage (FDR = 0.013), photosynthetic electron transport chain (FDR = 0.028), response to light stimulus (FDR = 0.044), and chloroplast localization. Detailed analyses (**Table 1**) showed three and six upregulated DEGs encoding core subunits of PSI and PSII, respectively. A significant upregulation was also found for genes related to photoprotective function, including Stress enhanced protein (SEP2) that binds to free chlorophyll (Ren *et al*. 2023), Early light-induced proteins (ELIP1, ELIP2), that prevent accumulation of free chlorophyll by the inhibition of its synthesis, and an ATP-dependent zinc metalloprotease (FTSH 8) involved in removal of damaged D1 protein. The C-induced upregulation was also found for the majority of identified plastid protein-coding genes, with 29 out of 59 genes showing increased expression. Interestingly, cold in both C and CLL plants upregulated CV (Protein CHLOROPLAST VESICULATION) responsible for stress-induced destabilization and degradation of chloroplasts (Wang & Blumwald 2015). However, the upregulation of genes related to photosynthetic apparatus was not found in CLL plants, and only two plastid encoded DEGs were upregulated. The subset of C-specific DEGs also included genes that reportedly play a role in stress tolerance and could correlate with the increased tolerance to freezing stress. These include down-regulation of transcription factor bHLH57, which was found to increase chilling tolerance in rice plants by activating trehalose synthesis (Zhang *et al*. 2023) and a downregulation of a gene encoding fatty acid elongase KCS12, At2g28630).

**Table 1.**
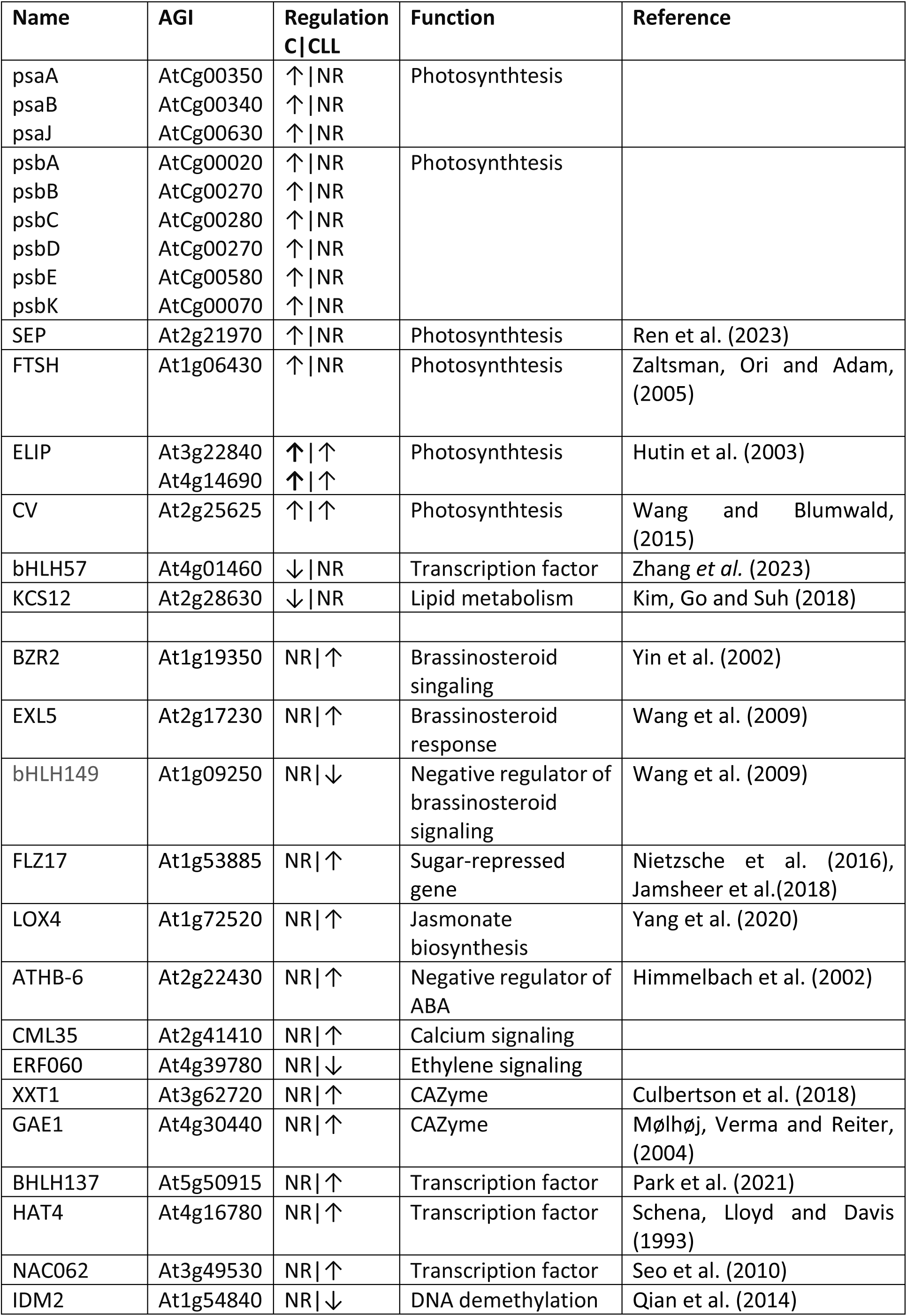
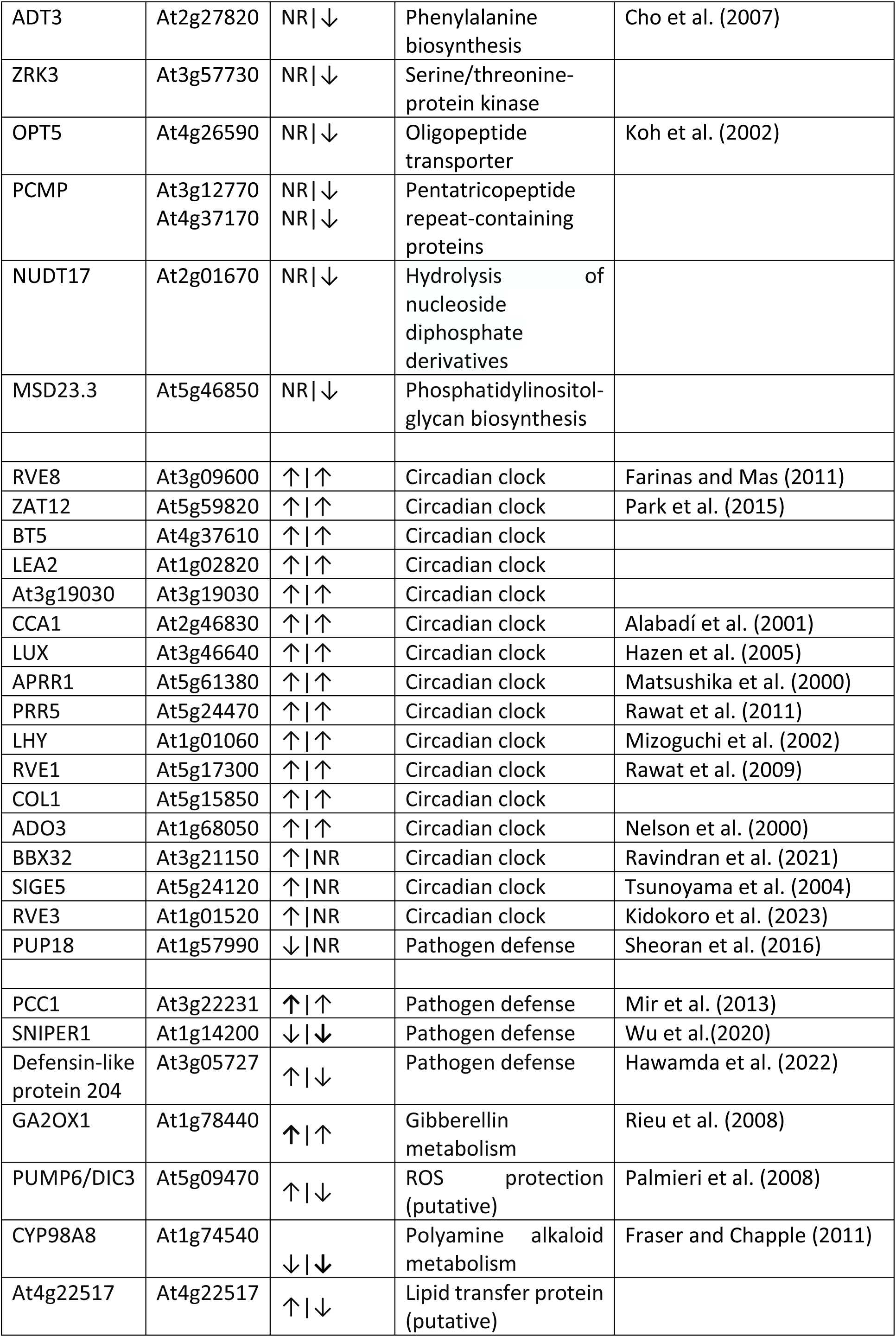
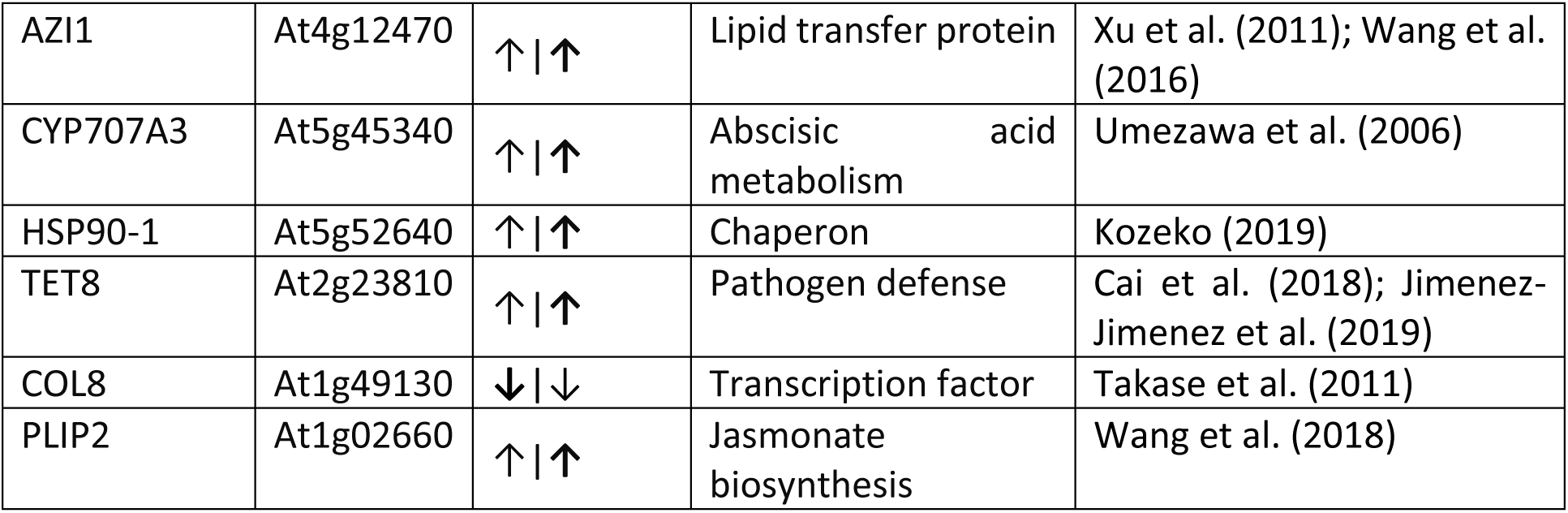
Differentially expressed genes with putative role in plant resilience to cold. Arrows mark regulations, and significant differences in C and CLL response are highlighted in bold. For details, see Supplementary Tables S1.

In total, 225 CLL-specific DEGs were identified **(Figure 3b)**. The upregulation of specific genes within the brassinosteroid signaling pathways highlighted its putative role in the CLL response, including *BRI1-suppressor 1* (BZR2) that exhibited a noteworthy fourfold increase, and *EXL5* (a role in a brassinosteroid-dependent regulation of growth and development). Additionally, *FLZ17*, identified as an interactor with SnRK1 (Nietzsche, Landgraf, Tohge & Börnke 2016; Jamsheer *et al*. 2018), was upregulated, suggesting its involvement in the direct control of phytohormone signaling under CLL. Moreover, the upregulation of *LOX4* and *ATHB-6* implies an active role in jasmonic acid biosynthesis and a negative regulation of ABA signaling pathway, respectively. Additional candidates of interests were *CML35* (potential calcium sensor), a transcription factor enhancing plant tolerance to pathogens by incorporation of cold mediated signaling *NAC062* (Seo *et al*. 2010), genes encoding carbohydrate metabolism enzymes xyloglucan 6-xylosyltransferase 1 (XXT1) and UDP-glucuronate 4-epimerase 1 (GAE1), Homeobox-leucine zipper protein HAT4, and transcription factor bHLH137. In total, 92 CLL-specific genes were downregulated. Genes with putative role in the CLL response included a negative regulator of the brassinosteroid signaling *bHLH149*, a regulator of DNA demethylation *IDM2*, *Nudix hydrolase 17* (NUDT17), *ADT3* (phenylalanine biosynthesis), *ZRK3* (Serine/threonine-protein kinase), *OPT5* (oligopeptide transporter), genes for pentatricopeptide repeat-containing proteins, *ERF060* (*Ethylene-responsive transcription factor*), and *MSD23.3* (phosphatidylinositol-glycan biosynthesis). Our analyses also pinpointed seven genes with previously unknown function, indicating their putative role in abiotic stress response (see **Supplementary Table S1** for details).

### 2.4 Circadian clock genes are part of early response to cold in young leaf

In *Arabidopsis,* the circadian clock is entrained by cold temperatures (Fowler, Cook & Thomashow 2005). Here, 59 DEGs associated with the circadian clock and rhythmicity were identified **(Supplementary Figure S1**). Approximately 45% of identified DEGs were observed in both the C and CLL plants. Five DEGs were found in all three treatments, namely *RVE8*, *ZAT12*, *BT5*, *LEA2*, and *AT3G19030* (**Table 1, Supplementary Tables 1**). The core clock genes, including *CCA1*, *LUX*, *APRR1*, *PRR5*, and *LHY*, were upregulated in both C and CLL plants. Furthermore, there was a noted upregulation of transcription factors linked to the circadian clock, such as *RVE1*, *COL1*, and *ADO3*. The transcription factor ZAT12 acts downstream of CBF1 and regulates cold acclimation and stress tolerance (Park *et al*. 2015; Zhang, Xiao, Wang, Khan & Liu 2024). The STRING analysis suggested a putative interaction between ZAT12 and the cold-upregulated circadian clock component LUX (**Figure 3d**). This interaction was also predicted by PEPPI (Bell, Schwartz, Freddolino & Zhang 2022), suggesting a previously unidentified direct link between the circadian clock and cold signaling pathways. More than 8% of circadian-responsive DEGs were exclusively observed only in C and LL plants, and the regulation was lost in CLL. Among the candidate DEGs of interest were the zinc finger transcription factor *BBX32* (integrates light and brassinosteroid signaling; Ravindran et al., 2021), circadian clock factor *SIGE5* (mediates expression of *psbD;* Tsunoyama et al., 2004), and *RVE3* (RVE family is involved in regulating expression under temperature stress; Kidokoro et al., 2023) which were upregulated in C, downregulated in LL, and not regulated in CLL plants (**Supplementary Figure S1**). In contrast, the transcript *PUP18* (belongs to the group of stress defense genes; Sheoran et al., 2016) encoding probable purine permease was up-and down-regulated in LL and C plants, respectively.

### 2.5 Low PPFD promoted cold stress response of genes involved in phytohormone metabolism and plant immunity

The comparative analyses of C and CLL plants revealed previously unidentified cold-responsive genes. Of particular interest were 100 DEGs that exhibited an enhanced cold stress response in CLL plants, showing an absolute fold change > 1.5 compared to C. The low-light promoted downregulation was found for DEGs encoding defense-related proteins PCC1 (regulates planťs pathogen defense through modulation of lipid content; Mir et al., 2013), SNIPER1 (E3 ligase involved in plant immunity response; Wu et al.,2020), Defensin-like protein 204, an enzyme of gibberellin catabolism GA2OX1, ELIP, Mitochondrial uncoupling protein 6 (PUMP6/DIC3), Cytochrome P450 98A8, and uncharacterized gene At4g22517. A promoted upregulation in CLL plants was identified for genes encoding lipid transfer protein (AZI1; participates in systemic acquired resistance and prevents electrolyte leakage in freezing conditions; Xu et al., 2011; Wang et al., 2016), an enzyme of abscisic acid catabolism Abscisic acid 8’-hydroxylase 3 (CYP707A3), HSP90-1 (chaperone involved in many processes, including R gene-mediated disease resistance), Tetraspanin-8 TET8 (reportedly involved in cell trafficking and plant immunity; Cai et al., 2018; Jimenez-Jimenez et al., 2019), a transcription factor COL8, FCS-like Zinc finger 13 (FLZ13, might facilitate interaction of SnRK complex with effector proteins; Nietzsche et al., 2016), and plastid phospholipase A1 PLIP2 that links ABA and jasmonic acid signaling (Wang et al., 2018).

### 2.6 The analyses of the early proteome response in young leaves indicated the significant involvement of ribosome composition and glutathione metabolism in the specific response observed in CLL plants

To complement transcriptomic analyses, the proteome of young leaf n. 6 (developmental stage L1.06) was analyzed. In total, the proteomic analyses provided identification and quantitative data for more than 5,000 and 2526 Arabidopsis protein families, respectively. A significant portion of the detectable proteome showed changes in response to cold or low light intensity. Differentially abundant proteins (DAPs; p<0.05; absolute fold change >1.4) represented 19.6%, 61.6%, and 60.2% of estimated protein content in C, CLL, and LL plants, respectively. However, the overlap in identified DAPs was formed by only 182 DAPs (**Figure 4a)**. The PCA separated the impact of light and cold in the first and second principal components, respectively. The proteome of CLL plants was clearly separated from that of S, C, and LL plants (**Figure 4b)**, and the two-way ANOVA confirmed an interaction between light and cold for more than 740 DAPs (**Supplementary Tables S2**). A substantial portion of DAPs in CLL plants showed a decrease in protein abundance compared to S. However, only carbohydrate-acting enzymes (CAZymes) and DNA metabolism exhibited a significant decrease (p<0.05), while the total protein content of major categories (31% for protein metabolism and 20% for photosynthesis) remained unaltered. Moreover, there was a notable increase in the total abundance of proteins related to secondary metabolism and components of RNA metabolism and processing (**Figure 4c-d**).

**Figure 4:**
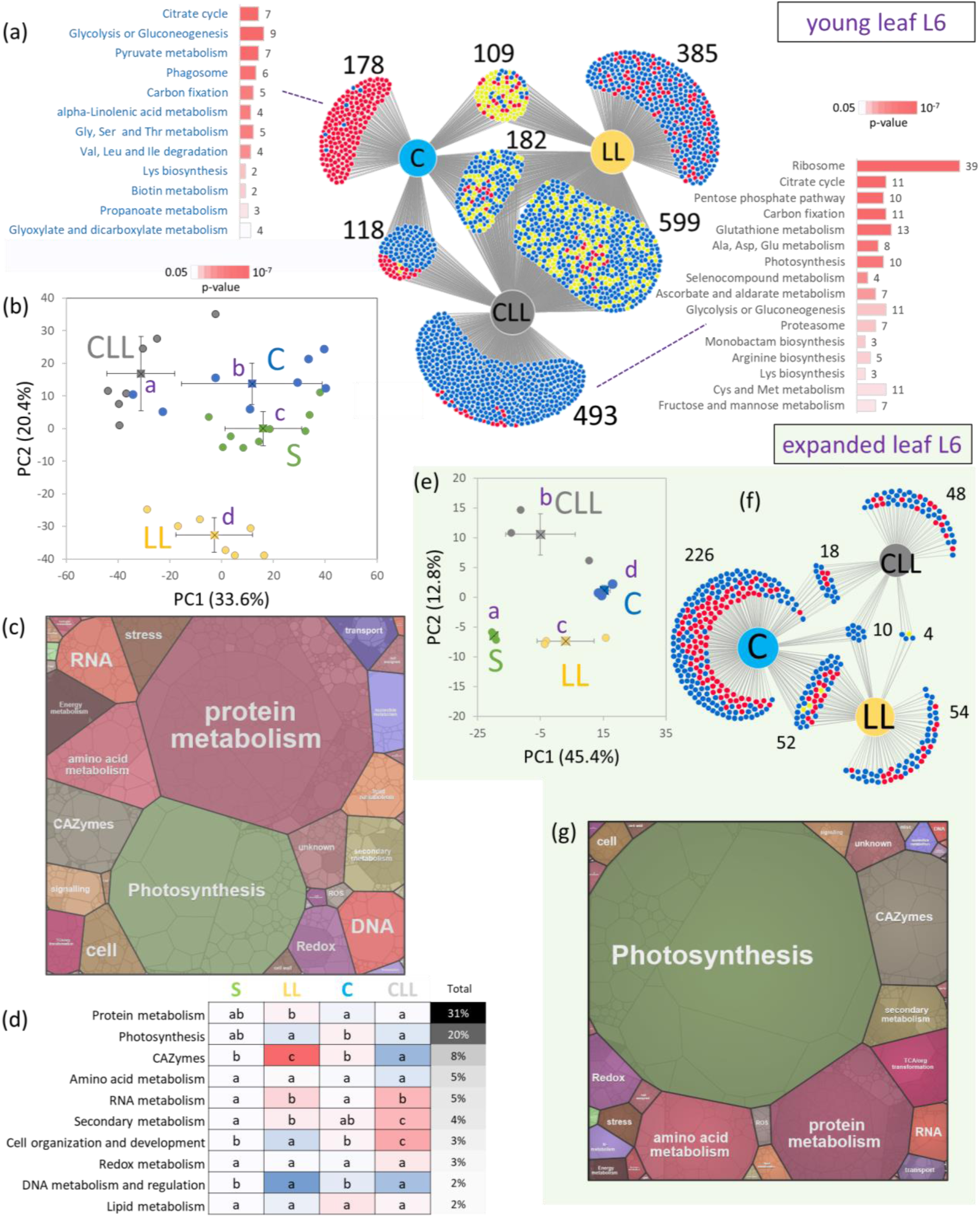
Cold response in leaf proteome. **(a)** DiVenn visualization of identified DAPs (p<0.05, absolute fold change >1.4) in young leaf L(1.06) and significantly enriched metabolic pathways identified by MetaboAnalyst in CLL and C specific DAPs; **(b)** PCA separation based on relative protein abundances of 2064 DAPs (p<0.05); **(c)** Proteome map illustrating the estimated protein content and the proportional distribution across diverse metabolic pathways within the young leaf of S plants, and **(d)** the corresponding comparison of ten most abundant categories. **(e)** The PCA separation based on relative protein abundances of 487 DAPs (p<0.05) identified in expanded leaf L(1.14); **(f)** The DiVenn visualization of identified DAPs (p<0.05, absolute fold change >1.4), L(1.14); **(g)** The proteome map of expanded leaf 6 of S plants. Circles in PCAs represent individual biological replicates, squares and lines represent group means and standard deviations, respectively; Letters indicate significant differences, Kruskal-Wallis and the Conover test, p<0.05); Red and blue dots in DiVenn indicate relative increase and decrease in protein abundances compared to S plants, respectively, while yellow dots represent differential responses between the comparisons. For details, see **Supplementary Tables S2**.

The detailed analysis of proteome data indicated a decrease in jasmonic acid metabolism in LL and CLL plants. Specifically, the abundances of allene oxide synthase CYP74A (a key enzyme in JA biosynthesis), two plastid-associated proteins PAPs (putative role in light/cold stress-related jasmonate biosynthesis), and UDP-glycosyltransferase UGT74D1 (glucosylates jasmonates) were significantly lower compared to S plants. Furthermore, cold-induced accumulation of lipoxygenase LOX2 (JA biosynthesis), was observed exclusively in C plants. CLL plants showed a decrease in abundances for enzymes of steroid biosynthesis (SMT2 and DIM), abscisic acid biosynthesis (zeaxanthin epoxidase ZEP), and auxin metabolism (Nitrilase NIT1, amidase AMI1). In contrast, polyamine biosynthesis seemed to be stimulated in all treatments (spermidine synthase SPDSYN2), and CLL plants displayed a remarkable reduction in amine oxidase CuAOalpha2, a polyamine degradation enzyme (**Table 2, Supplementary Tables S2**).

**Table 2.**
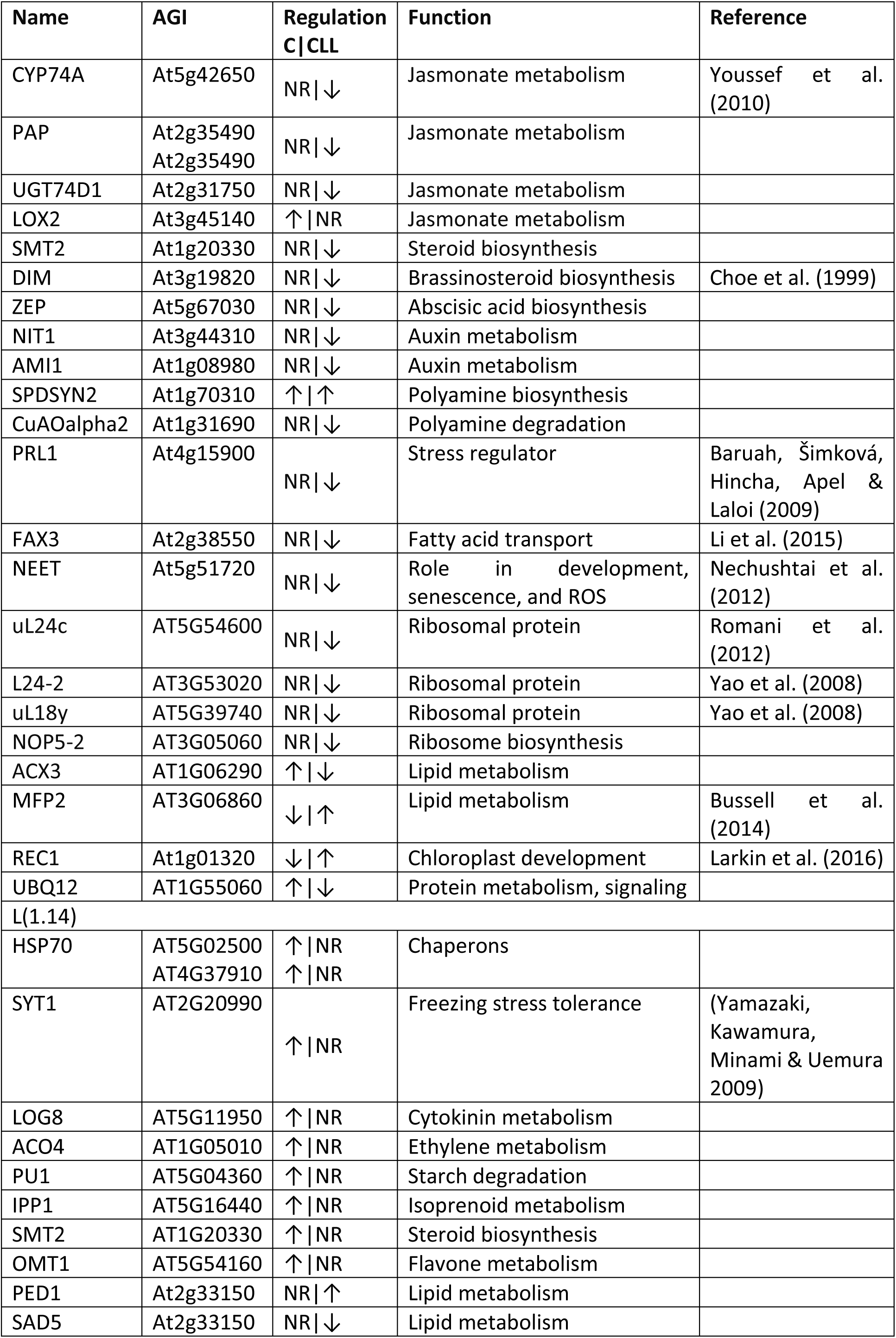
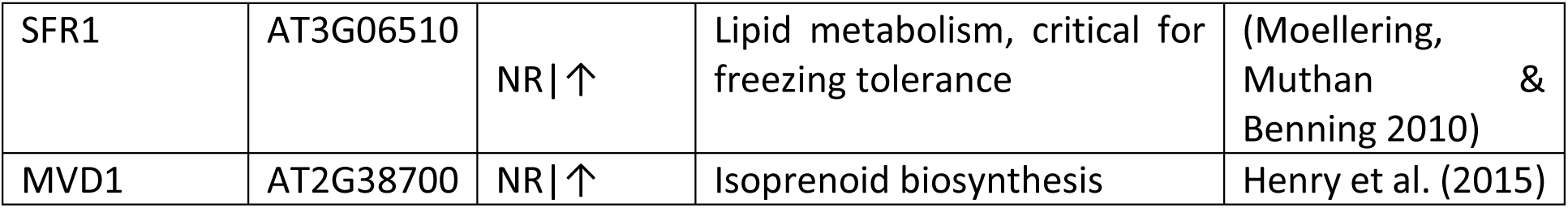
Differentially abundant proteins with putative role in plant resilience to cold. Arrows mark regulations. For details, see Supplementary Tables S2.

Proteins with a CLL-specific response with a putative role in cold stress included fatty acid transporter FAX3 and proteins involved in stress response and stress regulation (PRL1, NEET), and the analysis of metabolic pathway enrichment in CLL-specific DAPs showed an impact on citric acid cycle, glutathione metabolism, photosynthesis, CAZymes, amino acid metabolism, and ribosomes (**Figure 4a**, **Table 2**). The proteome of CLL plants exhibited a significant decrease in abundance of eukaryotic and chloroplast ribosomal proteins. Ribosomal protein content decreased in both C and CLL plants, with CLL showing a significantly more pronounced impact, decreasing on average by 28% compared to 11% in C. In total, 127 out of 187 quantified ribosomal proteins showed a significant decrease in abundance in CLL plants, including ribosomal proteins with documented significant impact on leaf growth and development uL24c, L24-2, and uL18y (Yao et al. 2008; Romani et al. 2012). The decrease was observed also for proteins involved in ribosome biosynthesis, including NOP5-2. For details, see **Table 2** and **Supplementary Table S2**.

The overlap between proteome response in C and CLL plants was substantial. In total, 301 DAPs were found in both datasets and only 11 showed contrasting response. Of particular interest were two peroxisomal enzymes involved in the fatty acid beta-oxidation pathway (ACX3, MFP2), protein REC1 that reportedly participates in chloroplast compartment size establishment (Larkin et al. 2016), and protein Polyubiquitin UBQ12 (**Table 2**).

### 2.6 Early proteome response in expanded leaf n. 6 identified CLL-specific alterations in lipid metabolism

To decipher the development-dependent mechanisms underlying cold stress response, the proteome of expanded leaf n. 6 (developmental stage L1.14) was subjected to proteome analysis and 2604 and 1982 protein families were identified and quantified, respectively. Similar to the young leaf, a distinct separation of CLL plants was observed (**Figure 4e**). In total, 412 DAPs were identified (p<0.05, absolute fold change > 1.4), including 228, 54, and 48 specific for C, LL, and C plants, respectively (**Figure 4f**, **Supplementary Table S2**). The majority of the expanded leaf n. 6 proteome represented proteins of photosynthesis, CAZymes, protein metabolism, and amino acid biosynthesis (**Figure 4g**). The results of the two-way ANOVA confirmed a notable interaction between light and cold conditions, specifically impacting 174 DAPs (adj. p<0.05; **Supplementary Table S2**). CLL plants exhibited the fewest number of DAPs, and the cold-induced responses observed in C plants were significantly weakened and completely abolished for 68 and 213 DAPs, respectively. The DAPs that were significantly accumulated only in C plants included two HSP70 chaperons, Synaptotagmin (SYT1, critical for maintaining plasma membrane integrity during freezing stress), hormone metabolism enzymes (LOG8 and ACC oxidase ACO4), Pullulanase PU1 (starch breakdown), and three enzymes of secondary metabolism pathways (IPP1, SMT2, OMT1). While the overlap in identified DAPs was limited (**Figure 4f**), metabolic pathway analysis revealed that the biosynthesis of secondary metabolites, amino acid metabolism, ribosomal proteins, glutathione metabolism, and CAZymes were found in all three datasets. The CLL plants showed significant enrichment in fatty acid metabolism, biosynthesis of unsaturated fatty acids, porphyrin and chlorophyll metabolism, 2-oxocarboxlic acid metabolism, and valine, leucine, and isoleucine degradation. Candidates of interest were enzymes of lipid metabolism (PED1, SAD5, SFR1) and isoprenoid biosynthesis (MVD1, mutant exhibits a significant decrease in campesterol and sitosterol content; Henry et al., 2015). Lastly, it’s worth mentioning that one of the presumed cold receptors and mediators of cold acclimation, the protein ANNEXIN 1, exhibited a 1.7-fold lower abundance in CLL plants compared to C or S. However, the regulation of this protein only marginally missed meeting the statistically significant threshold (p = 0.0524).

### 2.7 Comparison of transcriptome and proteome provided validation for the role of ROS metabolism, secondary metabolism, and lipids in CLL-specific response to chilling stress

A comparison of proteomics data from young leaves with NGS results revealed that 2447 proteins/genes were shared between the two methods, while 14102 transcripts were undetectable by the proteome analysis. Intriguingly, 75 quantified proteins were not captured by the transcriptomics dataset. A comparison of DAPs and DEGs revealed only 155 proteins/genes with overlapping profiles. Notably, only 16 and 49 displayed similar patterns in C and CLL plants, respectively. CLL-specific responses include components of ROS metabolism (catalase 3, L-ascorbate peroxidase 1, GSH transferases), protein At-NEET (a role in ROS metabolism and senescence; Nechushtai et al. 2012), and the aforementioned phytohormone metabolism enzymes (amine oxidase, UDP-glycosyltransferase UGT74D1, and auxin biosynthetic amidase). Apart from this, several enzymes involved in phenylpropanoid metabolism were also present in both datasets, alongside HSP70 proteins, Non-specific lipid transfer protein 2 (transfers phospholipids and galactolipids across membranes), and a key enzyme in the myo-inositol biosynthesis pathway IPS1. The complete list can be found in **Supplementary Tables S2.**

### 2.8 PLIP family is involved in cold acclimation and freezing stress tolerance

To affirm the hypothesized role of the promising candidates identified through the analyses, several Arabidopsis mutant lines were subjected to rigorous testing. The selection of mutant genotypes was based on availability, likelihood of observing functional impact from mutation, and expected gene/protein function. That encompassed mutants within *PLIP*, *FLZ13*, and *HSP90-1* where the cold-induced upregulation was significantly higher in CLL compared to C plants. The role of these genes in cold tolerance was assessed by exposing *Arabidopsis* mutant lines to cold stress as illustrated in **Figure 1a**. All three loss-of-function mutants demonstrated notably lower resilience to freezing stress, thus affirming their involvement in the cold response mechanism (**Figure 5a-c**). Omics data indicated a significant role of jasmonates in contrasting survival rates between C and CLL plants. Therefore, given the role of PLIP in jasmonate metabolism, the entire gene family comprising three members was analyzed in detail, and the impact on hormonome (**Figure 5d-e**, **Supplementary Tables S3**), jasmonate signaling (**Figure 5f-g**), survival rates (**Figure 5h**, **Figure 6a-b**), lipidome (**Figure 6c-g**), and chloroplasts (**Figure 7a-d**) was evaluated.

**Figure 5.**
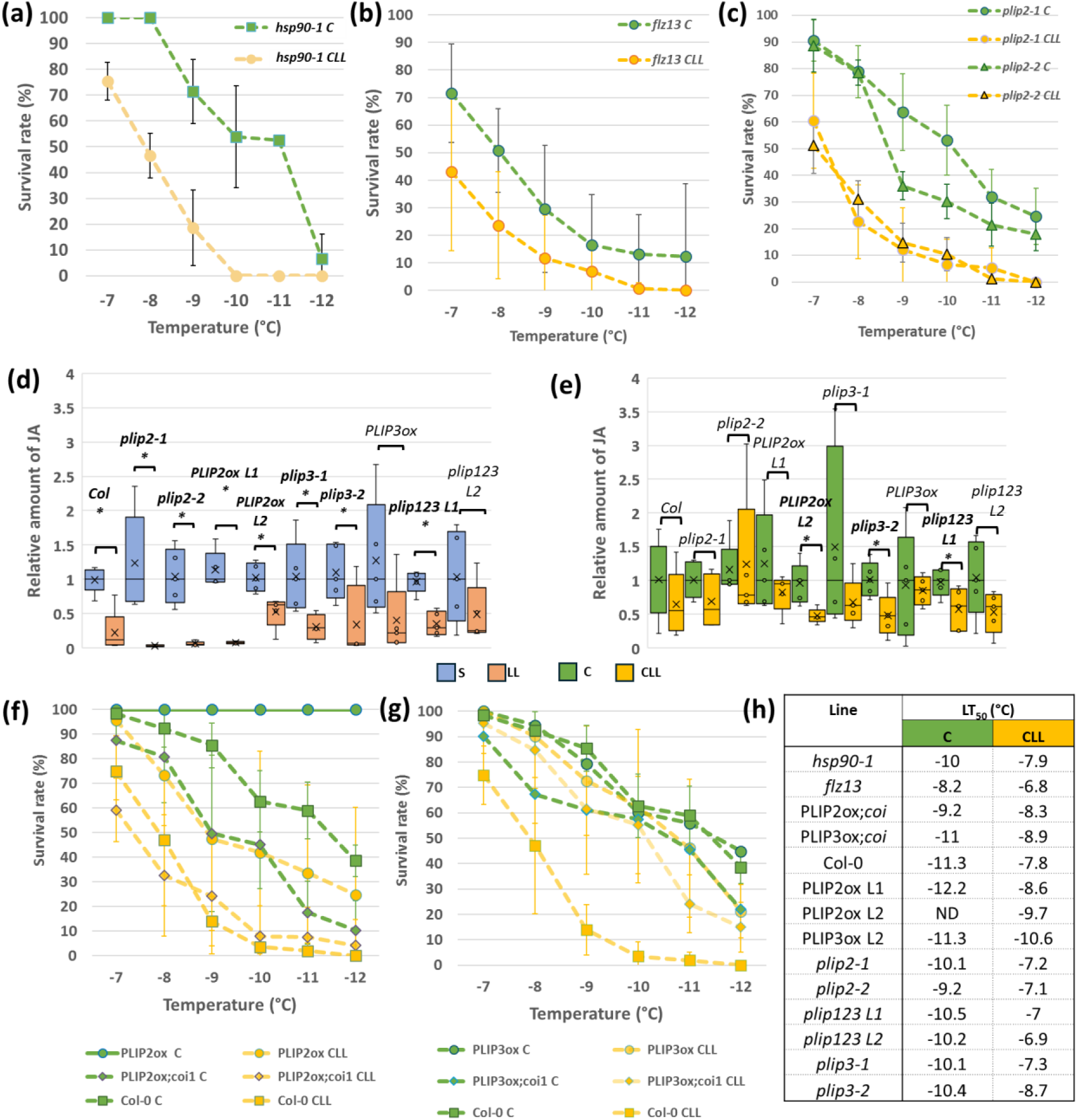
Mutations in candidate genes negatively impact freezing tolerance. **(a-c)** Survival rates of mutants in *HSP90*, *FLZ13, and PLIP*2. Means and standard deviation of at least three biological replicates (n=25); **(d-e)** Relative levels of jasmonate are predominantly regulated by light. This is a simplified comparison to highlight relative differences between S and LL plants, and C and CLL plants. Absolut values were normalized to respective S and C plants. The presented data represent results of five biological replicates, each pooled from at least 60 plants; **(f-g)** Modulation of JA perception affects freezing tolerance of plants overexpressing *PLIP2* and *PLIP3*. The presented data represent results of three biological replicates (n>50); **(h)** Calculated LT_50_ values for all mutants tested.

**Figure 6:**
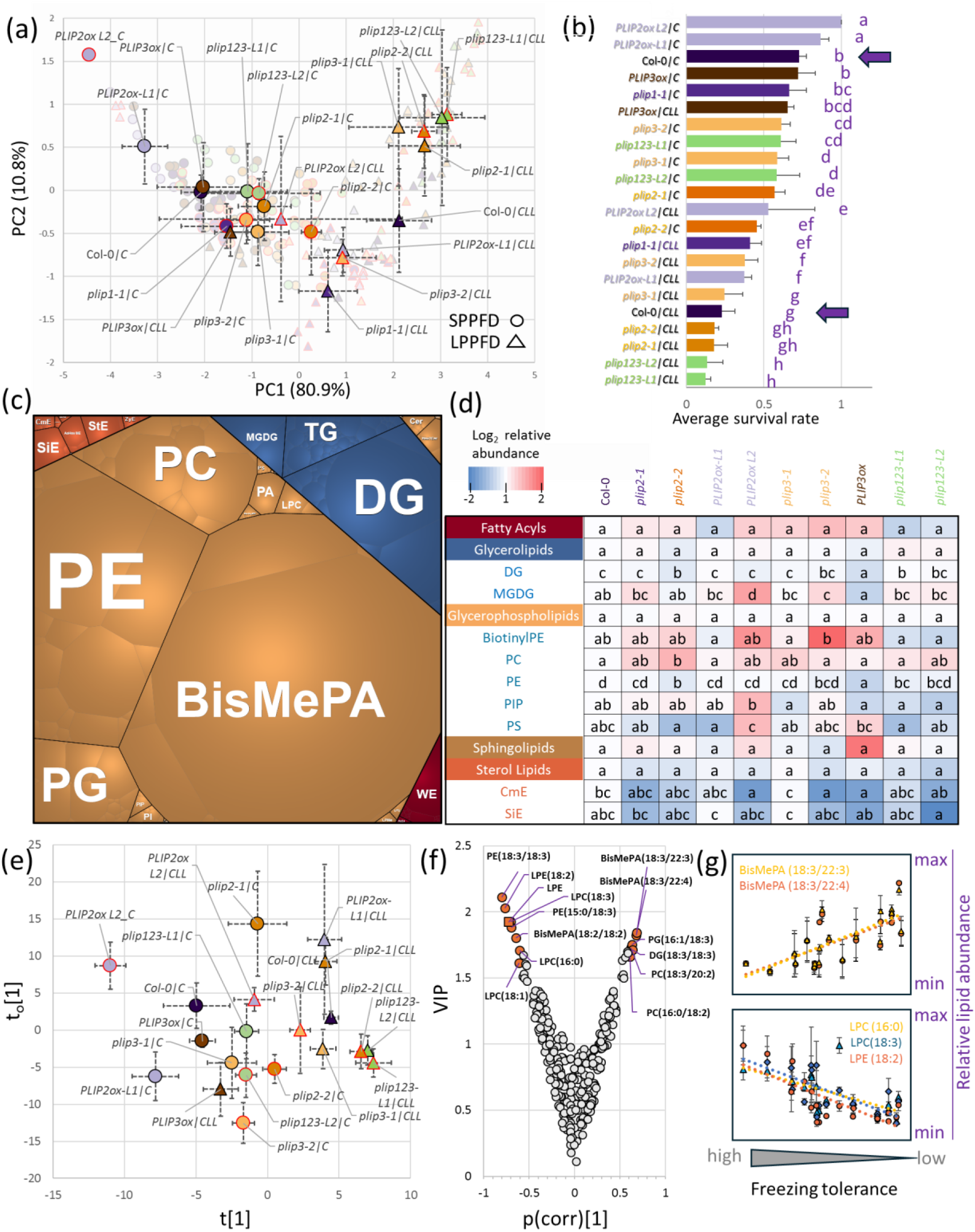
PLIP modulates lipidome and impacts freezing survival in a light-dependent manner. **(a)** PCA representation of survival assay data. Means and standard deviation of at least three biological replicates (n=25); **(b)** Mean survival rate in analyzed genotypes. Bars represent means and standard deviations, the letters represent significant differences (p < 0.05, Kruskal–Wallis and Conover’s test). Col-0 wild type plants are highlighted; **(c)** Average lipidome composition of Col-0 plantlets and **(d)** differences found in lipidome of the analyzed genotypes in control samples. Letters indicate statistically significant differences (ANOVA with Tukey’s HSD, p < 0.05); **(e)** Orthogonal partial least squares (OPLS) model based on freezing tolerance projection, the corresponding **(f)** Variable importance in projection (VIP) and identified lipid compounds correlating with freezing tolerance, and **(g)** selected examples of relative lipid abundance changes in response to freezing stress. Results are based on at least three biological replicates. For details, see **Supplementary Table S4**.

**Figure 7:**
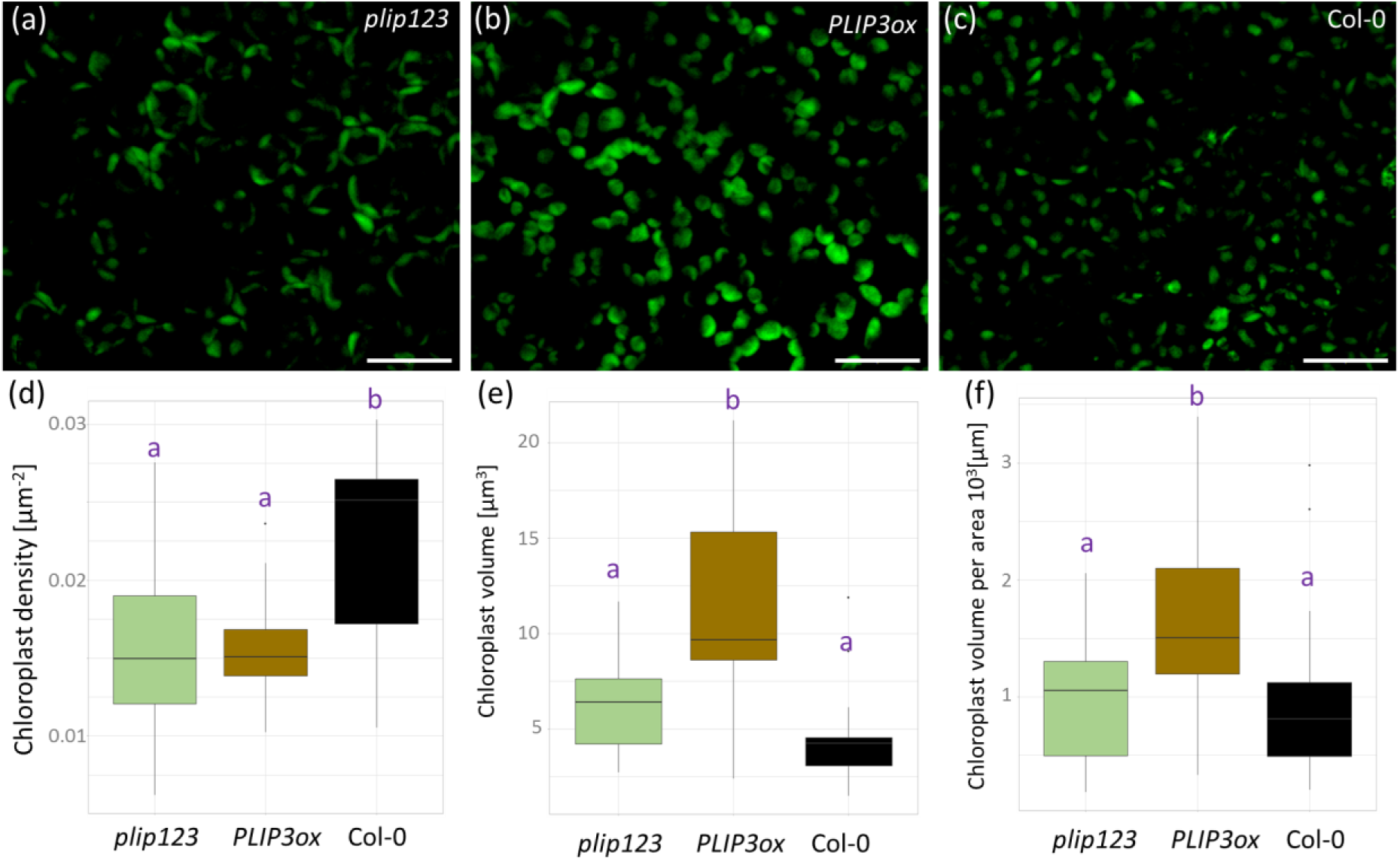
PLIP is a mediator influencing both the number and size of plastids. (a-c) Representative images of plastids in developing leaves of plip123, PLIP3ox, and Col-0 lines. Bar represents 20 µm; Box plot representations of (d) chloroplast density per area, (e) median chloroplast volume, and (f) median chloroplast volume per area. Results are based on three biological replicates each with five repeats. Letters in plots represent statistically significant differences (Kruskal-Wallis with Wilcoxon test, p < 0.05).

The experimental design for *PLIP* included available mutant and overexpressor lines, and the survival assay results were projected onto a PCA plot, distinctly segregating freezing stress tolerance along the first component. Moreover, it underscored a clear separation between plants that underwent acclimation under C or CLL conditions **(Figure 6a)**. The highest freezing resilience was identified in *PLIP2* overexpressor lines acclimated under C, followed by Col-0 and *PLIP3* overexpressor. Loss of function mutants *plip2*, *plip3* and *plip123* manifested significantly lower survival compared to Col-0 (**Figure 6b**). In general, plants acclimated under low PPFD had significantly lower survival rates compared to the corresponding lines acclimated under standard PPFD, with the lowest freezing stress tolerance identified in triple mutant. Remarkably, this decline was not observed in *PLIP3* overexpressors, which exhibited comparable levels of freezing tolerance in both C and CLL plants.

### 2.9 Jasmonate levels do not appear to be a downstream factor influencing PLIP2-mediated resistance to freezing

The constitutive overexpression of *PLIP2* and *PLIP3* reportedly triggers the excessive accumulation of bioactive forms of JA (Wang *et al*. 2018). The established correlation between this phytohormone and promoted plant resilience to low temperatures implied a potential link between JA levels and the observed resilience in *PLIP3ox* and *PLIP2ox* lines. To validate that, plants acclimated under S, LL, C, and CLL conditions for 147 hours were analyzed, monitoring JA levels alongside other stress-related phytohormones (**Figure 5d-e, Supplementary Tables S3**). The results showed a higher biological variability, but the JA pool was depleted in all LL plants compared to S, and most of these changes were statistically significant (p < 0.05). A comparable trend was noted when comparing CLL and C plants (**Figure 5d-e**), except for the *PLIP3ox* and *plip2-2* lines. The observed decrease in JA induced by low-PPFD is in line with proteomics results that indicated an attenuation in JA metabolism in LL and CLL plants, but it does not seem to correlate with the freezing stress resilience (**Figure 6a-b**).

SA levels were notably reduced in response to low-PPFD (**Supplementary Figure S2, Supplementary Tables S3**). The impact of cold mitigated this depletion, resulting in mostly insignificant differences between C and CLL plants. Interestingly, *PLIP3ox* and *PLIP2ox-2* exhibited increased SA levels in CLL plants compared to C. The difference did not reach statistical significance (p < 0.05), but it does correlate with the observed higher resilience to freezing stress. ABA was not impacted by low-PPFD under control temperature. However, a significant decrease in ABA was observed in CLL plants compared to C plants (**Supplementary Figure S2, Supplementary Tables S3**). While this regulation of the ABA pool doesn’t explain the enhanced resilience in *PLIP* overexpressors, it does underscore a distinct acclimation mechanism under low-PPFD.

The hormonome analysis did not show any pattern that would clearly correlate with the survival rates. However, the results of JA pool analysis could be misleading given the differences in the absolute JA content between different lines (**Supplementary Tables S3**). Next, the impact of *PLIP2* overexpression was assessed in lines that were crossed with mutant in a critical component of a receptor for jasmonates *CORONATINE INSENSITIVE1* (*PLIP2ox;coi1*; (Wang *et al*. 2018). The inhibition of JA signaling diminished the impact of *PLIP2* overexpression in both C a CLL plants (**Figure 5f-g**). Notably, *PLIP2ox;coi1* C plants exhibited freezing tolerance similar to that of *PLIP2ox* CLL, and *PLIP2ox;coi1* CLL had a similar LT_50_ to Col-0 CLL plants. That suggests that jasmonate signaling is integral to the PLIP2-mediated freezing resistance. Additionally, it potentially serves as a sensor of PPFD, aligning well with previous observations (Kazan & Manners 2011).

Interestingly, the analysis of *PLIP3ox;coi1* C did not reveal reduced resistance compared to *PLIP3ox* C plants. Additionally, the survival rates of *PLIP3ox;coi1* CLL were closer to those of *PLIP3ox* CLL than the wild type Col-0 (**Figure 5g-h**).

### 2.10 Shotgun lipidome profiling confirmed role of PLIPs in response to the combination of light intensity and highlighted putative lipid markers of freezing resistance

As PLIP is a lipid metabolism enzyme, lipidome profiles of individual lines were compared. The analysis of the leaf lipidome provided reliable identification and quantitation of more than 280 lipid compounds that represented more than 95% of the estimated lipid content. The leaf lipidome was formed predominantly by glycerophospholipids and glycerolipids, representing, on average, 77 and 18% of the Col-0 lipidome (**Figure 6c**). The comparison of lipid profiles clearly separated different genotypes with significant differences in abundance (ANOVA, p < 0.05) found for nine subclasses (**Figure 6d**). To eliminate biased evaluation of lipidome response to cold in individual genotypes, lipid abundances in response to cold were normalized to the respective controls. Next, the lipidome profiles of all genotypes were analyzed using the freezing tolerance projections (**Figure 6a**). The resulting orthogonal partial least squares (OPLS) modelling and the consecutive calculation of variable importance in projection (VIP) pinpointed a set of lipids that correlated with freezing tolerance: seven positively and six negatively (**Figure 6e-g**, **Supplementary Tables S4**).

### 2.11 Plastid number and size is impacted by PLIP

Next, three lines were selected for analysis of chloroplast size and quantity (**Figure 7a-f**), representing control (Col-0), line with the most attenuated resilience to cold (*plip123*), and line with a similar resilience in C and CLL plants (*PLIP3ox*). Col-0 displayed the highest number of chloroplasts per area (**Figure 7d**), but the median volume of these was lower than that in *plip123* or *PLIP3ox* (**Figure 6e**). The total volume of chloroplast per area was significantly higher in *PLIP3ox* (**Figure 6f**) and could represent the factor behind the observed improved resilience in *PLIP3ox* CLL plants.

## 3 DISCUSSION

### 3.1 PLIP plays an integral role in CLL-specific response to chilling stress

The primary objective of our analyses was to elucidate the contrasting molecular processes that underlie the acclimation responses of C and CLL plants. The complementary approaches based on transcriptome and proteome analyses both highlighted putative roles for lipid metabolism and transfer. Of particular interest was the significant upregulation of *PLIP2* (AT1G02660) that encodes plastidic lipase. While the abundance of this enzyme was below the detection limits of proteome profiling, its putative impact on jasmonates biosynthesis was corroborated by evidence from both proteome and transcriptome datasets. The Arabidopsis PLIP family comprises three members, PLIP1 (At3g61680) with a well-established role in seed oil biosynthesis, and PLIP2 and PLIP3, which are thylakoid-membrane-associated lipases that catalyze the initial step of jasmonic acid synthesis. Despite their shared function, PLIP2 preferentially utilizes monogalactosyldiacylglycerols (MGDG) as substrate, whereas PLIP3 prefers phosphatidylglycerol (PG) as its substrate (Wang et al., 2018). The experiment with mutant and overexpressor lines confirmed that PLIP2 abundance positively correlates with freezing stress resilience **(Figure 6a-b)**. The overexpressor lines displayed the highest survival rates, whereas the *plip2* mutant lines displayed the second lowest rates, marginally better than those observed in the triple mutant *plip123*. Moreover, the increased survival rates observed in *plip1* and *plip3* mutants compared to Col-0 in CLL plants might be associated with a potential compensatory mechanism and the accumulation of PLIP2 in these mutant variants. The experiment additionally validated that the absence of PLIP has a notably more pronounced impact under low PPFD conditions. While the highest decline in survival rates compared to Col-0 was 21% in C plants (*plip2*), it escalated to 10% in CLL plants (*plip123*).

### 3.2 The comparison of the proteome response between young and expanded leaves hints at a scenario where expanded leaves might be sacrificed to sustain the growth of young ones

As expected, the proteome profile of the expanded leaf 6 differed significantly from its earlier developmental stage at 1.06 **(Figure 4c,g**). There was a notable increase in the allocation of resources towards photosynthesis and CAZymes, while the processes associated with protein metabolism and RNA processing showed a distinct reduction. The comparison of quantified proteins revealed an overlap of 1514 proteins between both datasets, while 1012 were unique to dataset 1.06 and 469 to dataset 1.14 (**Supplementary Tables S2**). The response to cold was significantly attenuated in 1.14 plants, indicating that altering composition of these leaves was not a priority. The observed differences could correlate with the difference in plant size and the higher capacity to compensate stress response. However, it aligns with the established sink-source relationship, where actively growing leaves act as a sink for resources, while maintaining them comes at the expense of mature leaves (Chang and Zhu, 2017). Furthermore, it corroborates our findings that younger leaves exhibited greater resilience to cold stress compared to mature ones **(Figure 1c)**. Interestingly, 16 DAPs discovered in 1.14 CLL plants exhibited a similar trend to those identified in 1.06 CLL plants, including an enzyme of fatty acid biosynthesis (beta-ketoacyl-[acyl-carrier-protein] synthase III; AT1G62640, ↑), flavonoid biosynthesis (flavonol 7-O-rhamnosyltransferase; AT1G06000, ↑), a cis-acting element 14-3-3-like protein GF14 nu (AT3G02520, ↑), beta-amylase 3 (AT4G17090, ↓) that mediates accumulation of maltose upon freezing stress (Kaplan & Guy 2005), and chloroplastic protein WHY1 (AT1G14410, ↓) that maintains plastid genome stability (Maréchal *et al*. 2009).

### 3.3 Plastid maintenance could represent the major pathway that promotes freezing resilience in C plants

Major portion of DEGs and DAPs indicated significant alterations in photosynthesis and related metabolic pathways in C and CLL plants (**Figure 4A**, Table 1, Tables S1, S2). A notable upregulation of plastidic genes in C plants, might suggest that CLL plants are struggling to manage damage to their plastids. Alternatively, it could indicate that an inadequate PPFD isn’t providing enough energy to sustain fully functional photosystems. In a long term, that could explain lower resilience of CLL plants. That likely corresponds to the outcomes of the plastid volume analysis (**Figure 7a-f**). The observed lower quantity of chloroplasts in *plip123* mutant, in comparison to Col-0, seems to coincide with its status as the least resilient line. In parallel, *PLIP3ox*, the line displaying minimal PPFD impact on freezing resilience, demonstrated a notably larger total chloroplast volume than Col-0. It should be noted that plastids are also subject to degradation through autophagy, a process that serves both nutrient recycling and quality control purposes (Izumi *et al*. 2015). A notable rise in polyamine production in C and CLL plants (**Table S2**) might suggest NO-mediated autophagy (Minibayeva *et al*. 2023), where plants with a greater number/size of plastids could potentially benefit more.

### 3.4 Lipid composition correlates with freezing resilience

PLIP enzymes demonstrate a broad substrate specificity in vitro. Nevertheless, PLIP1 and PLIP3 exhibit a preference for cleaving chloroplastic phosphatidylglycerols, while PLIP2 primarily targets monogalactosyldiacylglycerol (MGDG) as its substrate (Wang, Froehlich, Zienkiewicz, Hersh & Benning 2017b). Beyond the previously documented influence on jasmonate biosynthesis attributed to PLIP2 and PLIP3, supported by our hormonal profiling (**Supplementary Table S3**), the impact of PLIPs extends to the modulation of lipidome composition (**Figure 6c-g**), and it is possible that this modulation could be the major reason for the observed freezing stress resilience. Lipid composition remodeling is crucial for cold tolerance (Gao *et al*. 2023) and here, the OPLS identified 13 compounds of interest that correlated with freezing resilience (**Figure 6f**), including *lyso*-phosphatidylethanolamines (LPE), *lyso-*phosphatidylcholines (LPC), and Bis-methyl phosphatidic acids (BisMePA). Both LPE and LPC (abundances positively correlate with resilience) were found in previous analyses of cold stress (e.g., Liu et al. 2022). Interestingly, LPE is an inhibitor of phospholipase D that mediates plant responses to stresses, and its inhibition promotes freezing tolerance of both non-acclimated and cold-acclimated plants (Rajashekar, Zhou, Zhang, Li & Wang 2006). The inhibition of phospholipase D additionally suppresses the production of phosphatidic acid, a trend that aligns with the observed negative correlation between BisMePA and both LPE levels and freezing resilience (**Figure 6g**).

## 4 MATERIALS AND METHODS

### 4.1 Plant material and growth conditions

To study the effect of light intensity on cold stress, we have employed the plant model *Arabidopsis thaliana* (L.) Col-0. Seeds were surface sterilized by immersion in 75% and 96% ethanol, respectively, and stratified in water for three days (4 °C, dark conditions). Plants were cultivated in AR-36L growth chambers (Percival Scientific Inc, Perry, IA, USA) under short-day photoperiod (65% relative humidity; 21/19 °C day/night temperatures, photon flux density (PPFD) 100 µmol.m^-2^.s^-1^ provided by fluorescent tubes Philips TL-D) using an Araponics hydroponic system (Araponics, Liege Belgium, 1.7 l tank) in a half-strength Murashige and Skoog media. Growth media was refreshed every seven days. Plants reaching the growth stages L(1.06) - (6 rosette leaves are greater than 1 mm) and L(1.14) were divided into four sub-groups (>42) and cultivated at the following conditions: (i) S-PPFD at (100 µmol.m^-2^.s^-1^), 21°C (S); (ii) low-PPFD (at 20 µmol.m^-2^.s^-1^; LL), 21 °C; (iii) S-PPFD at 4°C (C) and (iv) low-PPFD, 4°C (CLL). Ten leaves L(1.06), and expanded leaves L(1.06) from plants that just emerged developmental stage L(1.14) from three biological replicates were harvested after 3 hours of treatment (T3). Leaves were flash-frozen in liquid nitrogen, homogenized, and aliquoted for molecular analyses.

Plants for determination of lipid and hormone content of selected mutant lines mutant lines-*plip2-1*, *plip2-2*, *plip3-1*, *plip3-2, plip1,2,3* line 1, *plip1,2,3* line 2 and overexpression lines *PLIP2*-OX line 1, *PLIP2*-OX line 2 and *PLIP3*-OX line 2 (kindly provided by Prof. Christopher Benning, MSU) were prepared as follows: plants were cultivated on Petri plates for 2 weeks period at short day conditions (8h/16h light/dark). After this period, Petri plates were transferred into the following conditions:(i) S (ii) LL, (iii) C and (iv) CLL. The samples were collected for hormone and lipid analyses after T3 and 147 (T147) hour of exposure to the treatment. Plant material was snap-frozen in liquid nitrogen and stored at -80 °C until further use.

Validation assays for PLIP lines were performed as follows: *Arabidopsis thaliana* Col-0, and mutant lines *plip2-1*, *plip2-2*, *plip3-1*, *plip3-2, plip1,2,3* line 1, *plip1,2,3* line 2 and overexpression lines *PLIP2*-OX line 1, *PLIP2*-OX line 2 and *PLIP3*-OX line 2 (kindly provided by Prof. Christopher Benning, MSU) were cultivated on ½ Murashige-Skoog medium supplemented with 0.8% agar in 8/16 (day to night) day-length at S conditions. After two weeks of horizontal cultivation, plantlets were cold acclimated using C and CLL conditions for seven days. The freezing survival assay was performed using the adjusted method by(Hincha & Zuther 2020) with following changes: after a week of C-and CLL-acclimation, plants were exposed to decreasing temperature at rate 2°C per hour until the temperature in the growth chamber reached -6 °C. At this point the ice nucleation was induced. Next, plants were exposed to programmed cycle of temperature reduction -decrease of 1°C per 2 hours. Selected temperatures (-7°C, -8°C, -9°C, -10°C, -11°C and -12°C) were maintained for 2 hours, plates were harvested at the end of each maintenance period, Petri plates were removed and allowed to thaw at 4°C. After 14 days of recovery at S, the survival rates were calculated by counting the number of seedlings that produced new leaves.

For the chloroplast analysis, plantlets were cultivated at 21°C under S-PPFD for two weeks. The young developing leaves were observed with a confocal laser scanning microscope LSM700 (Carl Zeiss, Germany) equipped with a Plan-Apochromat 40× objective and an argon-neon laser with a wavelength 488 nm. Palisade mesophyll cells were scanned in five leaf regions with 448×362 frame (scaling per pixel: 0.2418 µm × 2.2418 µm × 0.25 µm). Image post-processing was performed using ZEN software (Carl Zeiss, Germany). Chloroplast volume and number of chloroplasts per area were determined using the Z-stacks and 3D Object Counter in ImageJ 1.54d (Schneider, Rasband & Eliceiri 2012). The experiment was done in three biological replicates.

### 4.2 RNA sequencing and differential expression analysis

The sequencing was performed at the Nucleomic Core facility (VIB, Leuven, Belgium, www.nucleomics.be). Samples were sequenced using single-end mode with a read 700 nt bp long using Illumina NextSeq500. Quality control was performed by using the Galaxy platform with FastQC; alignment was performed with the Salmon algorithm (Patro, Duggal, Love, Irizarry & Kingsford 2017). Sequences were searched against the Arabidopsis reference genome ARAPORT11. Differential gene expression (DEG) was determined for uniquely expressed genes with the EdgeR package (Robinson, McCarthy & Smyth 2009) against control treatment (S). Genes were determined to be differentially expressed with a false discovery rate (FDR) adjusted FDR ≤ 0.05.

Visualization and statistical analyses were performed with use or R packages: pheatmap (Kolde; 2015)., Bioconductor (Gentleman *et al*. 2004), ggplot2 (Wickham 2011), R software (v.4.0.3) (R Core Team, 2020, R Foundation for Statistical Computing, Vienna, Austria)

### 4.3 Proteomic analyses

Total protein extracts were prepared as described previously (Dufková, Berka, Psota, Brzobohatý & Černý 2023). Portions of samples corresponding to 5 µg of peptide were analyzed by nanoflow reverse-phase liquid chromatography-mass spectrometry using a 15 cm C18 Zorbax column (Agilent), a Dionex Ultimate 3000 RSLC nano-UPLC system and the Orbitrap Fusion Lumos Tribrid Mass Spectrometer (Thermo). Peptides were eluted with up to a 120-min, 4% to 40% acetonitrile gradient. Spectra were acquired using the default settings for peptide identification, employing HCD activation, resolution 60,000 (MS) and 15,000 (MS2), and 60 s dynamic exclusion. The measured spectra were recalibrated and searched against the Araport 11 protein database. Only proteins with at least two unique peptides were considered for the quantitative analysis. The quantitative differences were determined by Minora, employing precursor ion quantification followed by normalization. The mass spectrometry proteomics data have been deposited to the ProteomeXchange Consortium via the PRIDE (Perez-Riverol *et al*. 2022) partner repository with the dataset identifier PXD050271.

### 4.4 Lipidomic analyses

Total lipids were extracted as described previously (Dufková *et al*. 2023). In brief, 30 mg of plant material was extracted in terc-butyl-methylether:methanol mixture. The nonpolar fraction was separated and 200 µl aliquots were dried by vacuum centrifugation, resolved in 200 µl of isopropanol/methanol/ter-butyl-methyl-ether 4/2/1 supplemented with 20 mM ammonium formate and analysed by direct infusion using Triversa Nanomate (Advion Biosciences) nanoelectrospray source and the Orbitrap Fusion Lumos Mass Spectrometer (Thermo Scientific). Obtained spectra were analyzed by the software Freestyle 1.7 and LipidSearch 4.2 (Thermo Scientific).

### 4.5 Phytohormone analyses

Quantification of phytohormones and related compounds was performed using liquid chromatography-tandem mass spectrometry (LC-MS/MS) following the methodology by 10.1186/s13007-024-01165-8 Karady et al. (2024). cisOPDA was quantified according to Široká et al. (2022). For all compounds, concentrations were assessed using the standard isotope dilution method. All used solvents were of analytical or higher grade (Merck/Sigma-Aldrich KGaA, Darmstadt, Germany).

### 4.6 Statistics

Statistical tests were implemented using R, MetaboAnalyst (Pang *et al*. 2021), and the Real Statistics Resource Pack software for MS Excel (Release 6.8; Copyright 2013–2020; Charles Zaiontz; www.real-statistics.com). The reported statistical tests were generated and implemented using default and recommended settings unless otherwise indicated. Significant differences refer to p < 0.05 and adjusted adj. p < 0.05 (The Benjamini-Hochberg procedure, 5% FDR).

## ACKNOWLEDGEMENTS

We thank Dr. Lieven Sterck for his help with the RNA-sequencing analysis.

## CONFLICT OF INTEREST STATEMENT

The authors declare no conflict of interest.

## AUTHOR CONTRIBUTIONS

ML, DI, and JS initiated the study. ML and MČ designed the experiments. ML, Michaela Kameniarová (MK_1_), RK, SJ, ZQ, LT, and JS cultivated plants and collected plant material. ML, MD, and JS performed transcriptome analyses. MČ and ML performed proteome and lipidome analyses. ON and Michal Karady (MK_2_) performed hormone profiling. ML, MK_1_, KP, JN, JP, and LT performed validation assays. ML, MČ, and KP prepared figures. MD, ML, MČ, DI, and JN contributed to data analysis and interpretation. ML and MČ wrote the draft. MD, JN, and DI reviewed the manuscript. ML and MČ finalized the manuscript. All authors approved the published version.

## FUNDING

Funding for this work was provided by the Czech Science Foundation (grant number 20-26232S) and the Ministry of Education, Youth and Sports of the Czech Republic with support from the European Regional Development Fund (grant no. CZ.02.1.01/0.0/0.0/16_019/0000738, project name “Centre for Experimental Plant Biology”).

## DATA AVAILABILITY STATEMENT

The data that support the findings of this study are available in supporting information and in the following data repository: https://www.ncbi.nlm.nih.gov/geo/, accession GSE278942; http://www.ebi.ac.uk/pride, accession PXD050271.

## Supporting Information

Supplementary Table S1. Supporting data for transcriptomics. Supplementary Table S2. Supporting data for proteomics.

Supplementary Table S3. Supporting data for hormonomics. Supplementary Table S4. Supporting data for lipidomics.

Supplementary Figure S1. Regulation of circadian responsive genes. Supplementary Figure S2. Phytohormone analyses.

## Notes

### Competing Interest Statement

The authors have declared no competing interest.

