## Supplementary material for "Light quantity impacts early response to cold and cold acclimation in young leaves of Arabidopsis": Supplementary Figures.docx

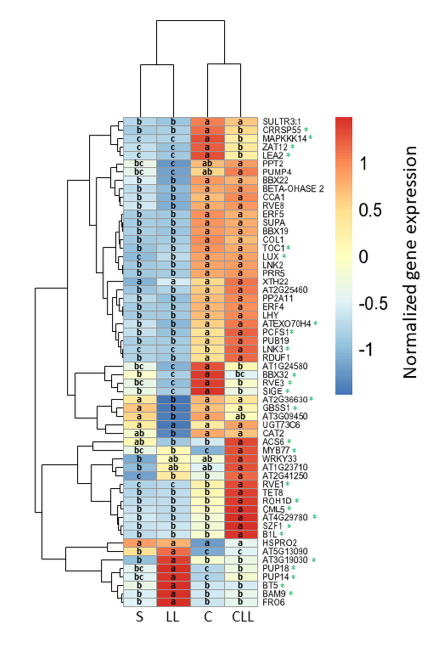

**Figure S1:** Combination of C- and low-PPFD impacted regulation of circadian responsive genes. DEGs found in RNA-seq analyses after 3-hour treatment. Significant interactions between low-PPFD and cold (p-value<0.05) are highlighted with an asterisk. The results of two-way ANOVA and Tukey post hoc test (p-value<0.05). For details, see Supplementary Tables S1.

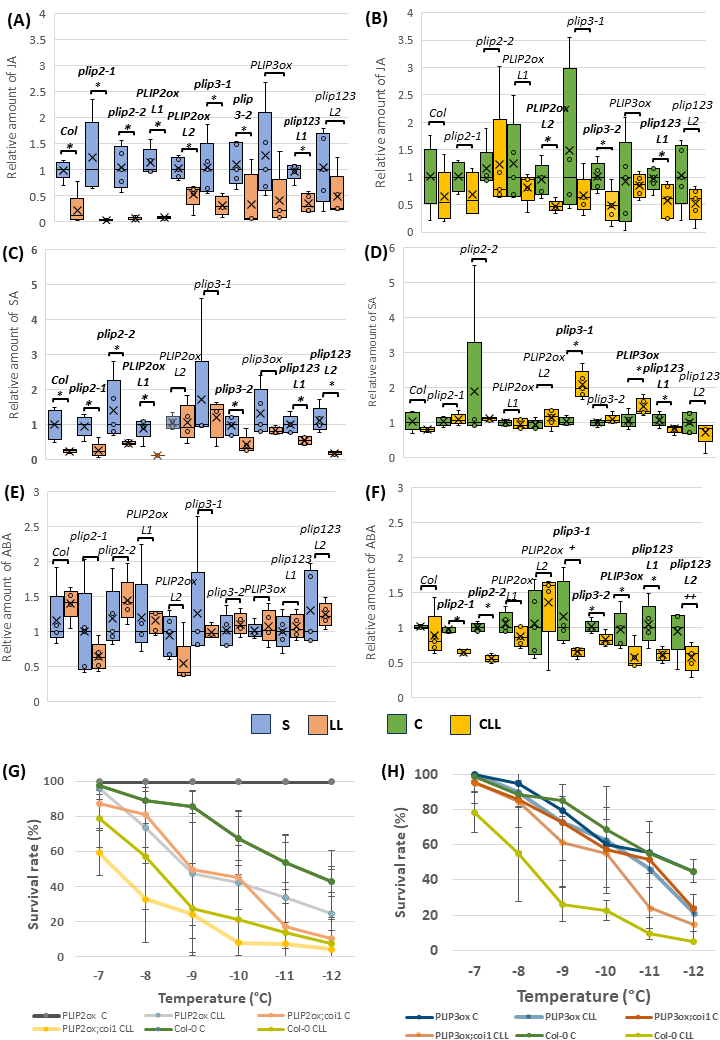
**Figure S2**. **Relative levels of plant hormones are predominantly regulated by light.** A simplified comparison to highlight relative differences between S and LL plants (A, C, E), and C and CLL plants (B, D, F). Absolut values were normalized to respective S and C plants. The experiment design and abbreviations are identical to those in Figure 6A. The presented data represent results of five biological replicates, each pooled from at least 60 plants. For details, see Supplementary Tables S3.
